# Capturing Protein-Ligand Recognition Pathways in Coarse-grained Simulation

**DOI:** 10.1101/868687

**Authors:** Bhupendra R. Dandekar, Jagannath Mondal

## Abstract

Protein-substrate recognition is highly dynamic and complex process in nature. A key approach in deciphering the mechanism underlying the recognition process is to capture the kinetic process of substrate in its act of binding to its designated protein cavity. Towards this end, microsecond long atomistic molecular dynamics (MD) simulation has recently emerged as a popular method of choice, due its ability to record these events at high spatial and temporal resolution. However, success in this approach comes at an exorbitant computational cost. Here we demonstrate that coarse grained models of protein, when systematically optimised to maintain its tertiary fold, can capture the complete process of spontaneous protein-ligand binding from bulk media to cavity, within orders of magnitude shorter wall clock time compared to that of all-atom MD simulations. The simulated and crystallographic binding pose are in excellent agreement. We find that the exhaustive sampling of ligand exploration in protein and solvent, harnessed by coarse-grained simulation at a frugal computational cost, in combination with Markov state modelling, leads to clearer mechanistic insights and discovery of novel recognition pathways. The result is successfully validated against three popular protein-ligand systems. Overall, the approach provides an affordable and attractive alternative of all-atom simulation and promises a way-forward for replacing traditional docking based small molecule discovery by high-throughput coarse-grained simulation for searching potential binding site and allosteric sites. This also provides practical avenues for first-hand exploration of bio-molecular recognition processes in large-scale biological systems, otherwise inaccessible in all-atom simulations.

## Introduction

Protein-ligand binding involves complex recognition process. ^1^ Precise knowledge of the underlying dynamical recognition pathway can potentially pave the way for direct elucidation of underlying mechanism.^2^ Towards this end, all-atom molecular dynamics (MD) simulation has lately emerged as the method of choice, thanks to its proven ability to capture the act of the ligand spontaneously binding to its designated protein cavity at atomic resolution and in real time.^3,4^ In this communication, we demonstrate that, coarse-grained models,^5,6^ when systematically improvised, can offer an attractive and inexpensive alternative of all-atom simulation towards complete capture of protein-ligand binding events, from solvent to protein cavity. This is achieved within a significantly shorter wall clock time compared to that of all-atom MD simulations.

Over the last few years, multiple independent investigations^3,4,7–11^ on complex systems have demonstrated that all atom molecular dynamics simulations can predict the complete biomolecular recognition process in multi-microsecond long time scale, where the ligand, initially placed in bulk solvent, spontaneously diffuses through the solvent to the protein and eventually finds and binds its designated protein cavity. The trajectories of these all atom simulations have remained very useful in revealing crucial rate-limiting processes, namely transient helix-gate opening^8^ or ligand-induced cavity opening,^9^ which act as precursor to the eventual binding event. However, these all-atom MD simulations of protein-ligand recognition processes are always associated with a prohibitively large computational expenses, because of the large system sizes and requirements of long time scale simulations. More over, the stochastic nature of these processes also requires one to perform these time-intensive processes multiple times, which adds to the overall computational expense. While there has been recent surge in the usage of adaptively sampled short time scale trajectories to infer the long time scale binding event within the framework of Markov state models, ^10,12^ the requirement of the large number of simulation trajectories (towards realising Markovian criterion) has not helped in reducing the computational burden. This prompted us to explore the possibility of a practical alternative which can capture the full process of protein-ligand binding at a reasonable resolution but at the cost of a frugal computational expense.

On the other hand, a reduced representation of the biologically relevant system and solvent that retains the essential molecular aspects for the system of interest, enables faster sampling due to reduced degrees of freedom. In this spirit, coarse-grained (CG) models of biomolecules have been around the corner for a while now. ^5,6^ The usage of CG models in a variety of simulation techniques has proven to be a powerful tool to probe the spatial and temporal evolution of systems, beyond what is feasible with traditional all-atom models. This has seen the application of coarse-grained models in exploring multiple disciplines of biological processes: exploring lipid membrane properties, ^13,14^ protein-lipid interactions and antimicrobial activities,^15,16^ membrane protein oligomerization,^17,18^ surfactant,^19^ peptide and protein self-assembly,^20–22^ membrane fusion,^23,24^ structural dynamics of polymers etc. Recently, coarse-grained models are also being extended to study ribosome dynamics ^25^ and various protein-DNA transient complexes.^26–28^ Motivated by these success stories of coarse-grained models, we speculated if coarse-grained models of proteins and small molecules can be put into play to substitute computationally expensive all-atom simulations for capturing protein-ligand recognition process on the fly. In this work, by building on popular MARTINI^6,29–31^ coarse-grained frame-work, we report that while its vanilla implementation on protein-ligand system might fail to capture the binding event, judicious optimization of elastic network^32^ in MARTINI protein model enable the simulation of the spontaneous binding process of ligand to the designated cavity at crystallographically identical pose.

## Results and Discussion

In this work we have considered three prototypical protein-ligand systems as shown in Figure 1, namely : a) trypsin recognizing benzamidine as a ligand, b) benzene binding to L99A mutant of T4 Lysozyme and c) camphor binding to cytochrome P450 in its heme active site. All three systems have remained the subject of prior experimental investigations ^33–35^ and all-atom simulations^7–9^ and hence would serve as suitable benchmark to verify the ability of the coarse-grained simulations to mimic the recognition process. More over, these three systems also offer a set of diversity in their traits of recognition process: Trypsin/benzamidine system presents a solvent-accessible ligand binding site,^7,33^ while cytochrome P450/camphor ^9,34^ system demonstrates a deeply buried or solvent-inaccessible active site for substrate recognition. Finally, the process of benzene recognising a cavity in L99A mutant T4 Lysozyme has recently been found to follow multiple pathways.^8,36^ All these diversities in three system pose suitable challenges for a coarse-grained approach to succeed.

**Figure 1:**
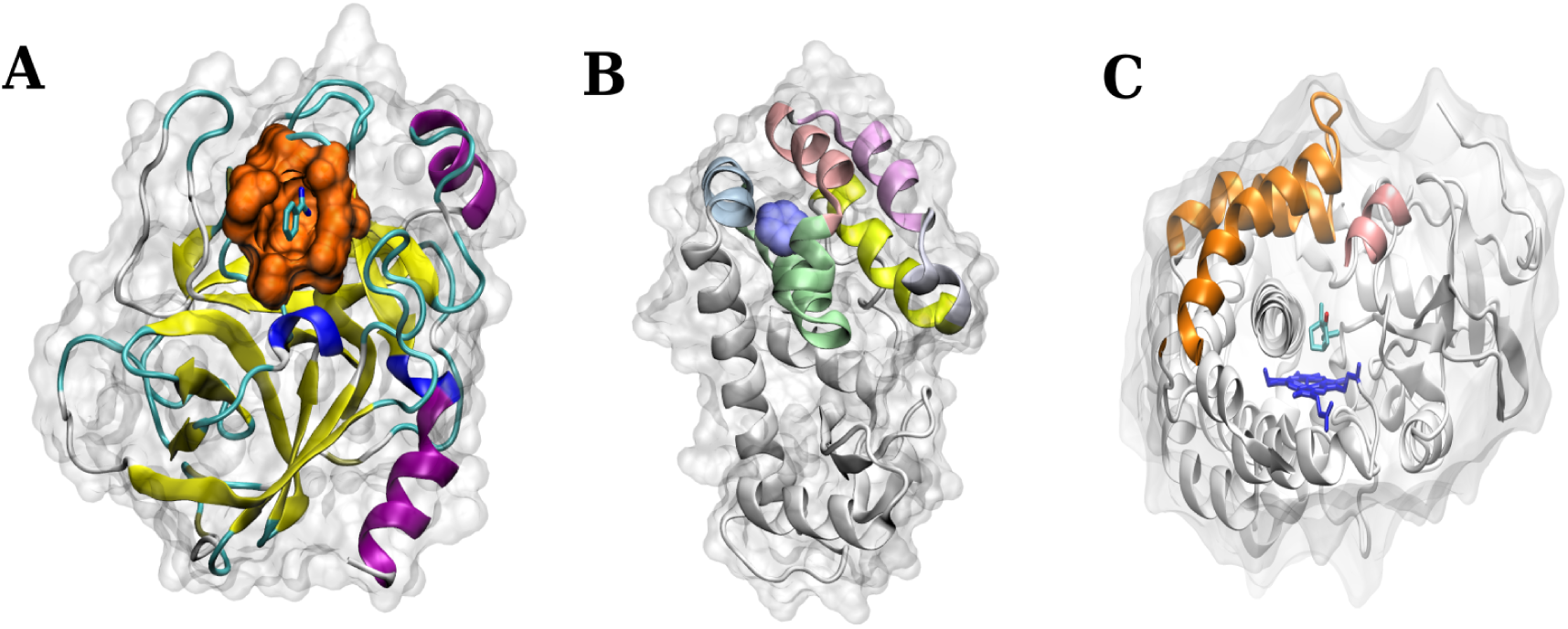
Protein ligand binding systems under study. (A) Trypsin and Benzamidine (PDB:3ATL) with solvent accessible cavity. (B) L99A T4lysozyme and Benzene (PDB:3DMX) with buried cavity. (C) Cytochrome P450 and Camphor (PDB:2CPP) with buried cavity.

We first attempted routine implementation of MARTINI mapping of these three proteins. The coarse-grained parameters for the ligand molecule are also individually optimised by fitting bonded interactions with all-atom simulation and using analogous MARTINI atom-types(see figure S1 and S2 and methods in SI). On the other hand, all-atom protein model is first coarse-grained using 4:1 mapping and the parameters of bond-connectivities are imposed as per the secondary structure of the protein sequence and MARTINI atom-types. ^30^ However, all our initial attempts in simulating the ligand diffusion through the protein and solvent and eventually capturing the act of ligand getting to the cavity were futile, even though the ligand came closer to the protein cavity at a number of occasions. A closer inspection of these unsuccessful trajectories revealed that the MARTINI model, in its regular implementation, fails to maintain the three-dimensional fold or tertiary structure of the protein. More importantly, as represented in movie S1 of apo protein of trypsin and in the time profile of root mean squared deviation (RMSD) from the crystal structure (figure S3), the binding cavity of trypsin, in its coarse-grained description, occasionally gets occluded and fails to accommodate the incoming ligand.

The observation that the regular implementation of the MARTINI coarse-grained approach does not allow for capturing protein-ligand binding process, implied that an optimisation of MARTINI framework is necessary so as to maintain the tertiary fold of the protein. Accordingly, we introduced elastic network among the backbone beads of the protein model, in accordance with Periole and coworkers ^32^ and subsequently optimised the coarse-grained model. Specifically, we optimised two parameters pertaining to elastic network, namely force constant (*K*_*f*_) and cut-off length (*R*_*c*_) (see figure 2) via iterative comparison with RMSD of all-atom model of the apo form of the proteins. Figure 2 B-C shows that a combination of *R*_*c*_=0.9 nm and *K*_*f*_ =500 kJ/mol/nm^2^ furnishes a reasonable match in the protein RMSD between the all-atom and coarse-grained model for trypsin. The root mean squared fluctuations (RMSF) of the residues of the apo-proteins obtained from all-atom models were also found to be in agreement with the coarse-grained model with optimised elastic networks (see figure 2D).

**Figure 2:**
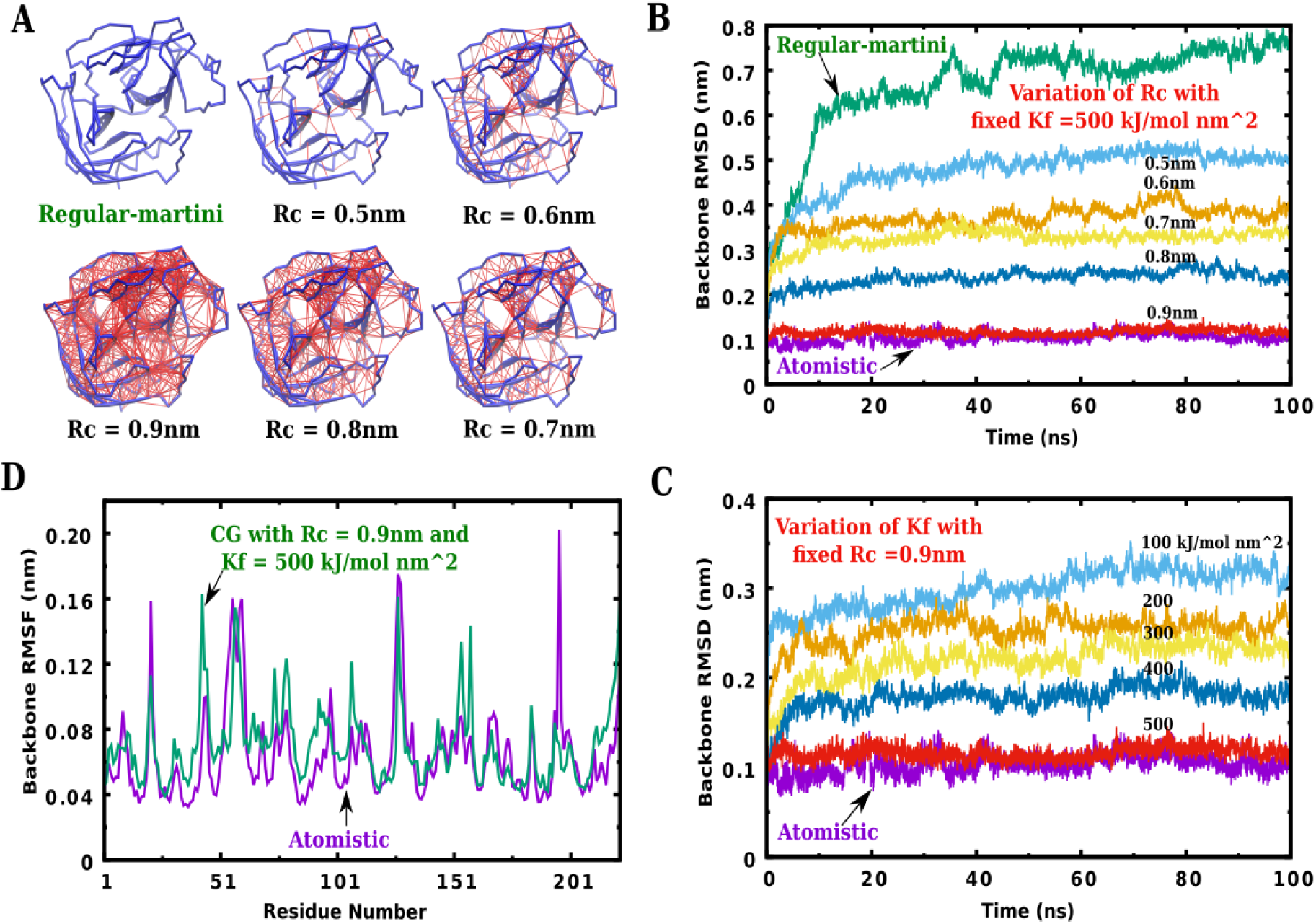
Optimization of elastic network parameters Rc and Kf for trypsin CG protein. (A) elastic network applied on the backbone (shown in blue licorice representation) of CG trypsin protein and the network is shown in red lines. (B) Comparision of CG backbone RMSD with atomistic RMSD for various cut-off (*R*_*c*_) distance of same *K*_*f*_ and (C) for different force constant (*K*_*f*_) of same *R*_*c*_. (D) Comparison of Root mean squared fluctuations (RMSF) of all protein residues obtained via all-atom model and coarse-grained models with optimised parameter of elastic networks

Subsequently, we modelled trypsin using elastic network corresponding to these optimised parameters and simulated the diffusion of benzamidine molecule (separately coarse-grained, as described in SI method and figure S1) in the aqueous solution of the trypsin. SI Movie S2 demonstrates a representative simulation trajectory of trypsin-benzamidine recognition process in the coarse-grained framework. We find that benzamidine, starting from the bulk, diffused through the solution and protein surface and eventually identified the solvent-accessible binding site and bound there efficiently. As compared across figure 3A-C, the final simulated bound pose, as obtained from the current coarse-grained model is in excellent agreement with the crystallographic pose (pdb 3ATL). ^37^ From the time profile of the cavity-benzamidine distance and benzamidine RMSD (relative to crystallographic pose), (Figure 3D-E), the eventual match of the bound pose with the crystal structure is evident. The trypsin-benzamidine binding phenomena were found to be reproducible across multiple independent trajectories. Overall, we find the coarse-grained simulation successfully captures the trypsin-benzamidine binding process, and most importantly at the cost of a meagre computational expense compared to all-atom simulation.

**Figure 3:**
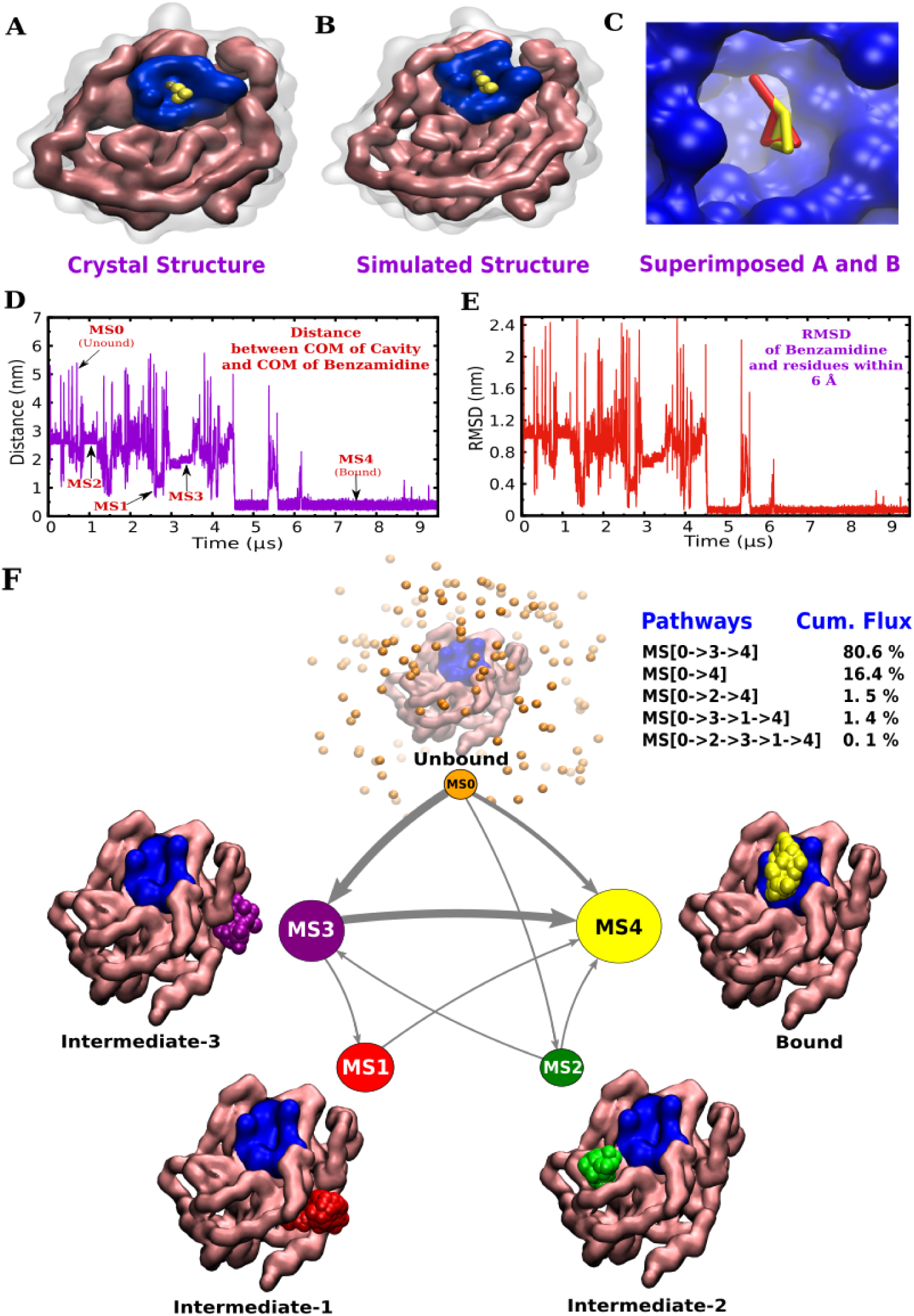
(A) The crystal structure of CG-3ATL. (B) The simulated structure of CG-3ATL. The protein structures are shown in quick surf representation, for clearity only backbone(mauve) and cavity(blue) is shown in opaque colors rest of the protein is kept transparant. The CG-benzamindine is shown in vdw representation. Protein heavy atoms within 6 *Å* of benzamidine in the CG-3ATL are defined as the “binding cavity”. (C) Close view of benzamidine pose after superimposing backbones of crystal structure((A) with red-benzamidine) with simulated structure(B with yellow-benzamindine). (D) The time profile of the distance between COM of cavity and COM of benzamidine and (E) The time profile of RMSD of benzamidine with residues within 6 Å (relative to crystallographic pose). (F) The five state Markov State Model(MSM) for trypsin benzamidine binding. For clarity only polar bead-type (P4) of coarse grain benzamidine is shown in the macrostates. The kinetic 7 network between these five state is obtained form Tansition Path Theory (TPT) calculations. The thickness of the arrow is proportional to net flux going from unbound state MS0 to Bound state MS4. The table on the top right corner of the figure shows the cumulative fluxes for each of the five benzamidine binding pathways.

The computationally affordable simulation of efficient ligand recognition process, via coarse-grained approach in combination with optimized elastic network, prompted us to explore the possibilities of enumerating all binding mechanistic pathways in a more quantitative and exhaustive fashion. Accordingly, we spawned a large set of multi-microsecond long binding simulation trajectories (see table 1) and additionally adaptively sampled numerous short-length simulation trajectories (total 128 microsecond) of trypsin-benzamidine interaction, within the coarse-grained model, and developed a Markov-state model (see method in SI) (MSM).^38–40^ The implied time scale levelled off beyond a lag time of 20 ns (figure S4B) and the model also successfully passed Chapman-Kolmogorov test,(figure S4C) which allowed us to build a converged MSM using the simulation trajectories of trypsin-benzamidine.

**Table 1:** Summary for protein-ligand Binding simulations using coarse-grained model

| Protein-Ligand System | No. of 10μs Simulations | No. of Bound Trajectories | No. of Binding Pathways | Simulated Binding Rates<br>$M^{-1}s^{-1}$ | Experimental Binding Rates<sup>33,44-46</sup><br>$M^{-1}s^{-1}$ |
| --- | --- | --- | --- | --- | --- |
| Trypsin-Benzamidine | 30 | 22 | 5(from TPT) | $3.68 \times 10^8$ | $0.29 \times 10^8$ |
| T4Lysozyme-Benzene | 30 | 17 | 5 | $8.22 \times 10^6$ | $(0.8 - 1) \times 10^6$ |
| CYP450-Camphor | 30 | 6 | 2 | $0.34 \times 10^7$ | $2.5 \times 10^7$ |

The key macro states, as obtained via subsequent lumping of micro states (see methods in SI) are able to recover five important ligand locations around protein: a macro state where the ligand is unbound MS0, a macro state where the ligand is completely bound to pocket of trypsin MS4 and additionally three interesting intermediates MS1, MS2, MS3 as macro states where the ligand is located in different parts of the trypsin. Interestingly all five macro states obtained in the CG binding simulations are similar to key binding spots reported in the earlier atomistic simulations done by Fabraitis and coworkers^7^ and more recently by Noe and coworkers. ^10^ These macrostates are characterised in the table S3. We obtained the kinetic network of transition between the metastable states using transition path theory. The kinetic network connecting all of the five meta stable states suggests that there are possibilities of multiple binding pathways. (See Figure 3F) Interestingly the current coarse-grained simulations recover the the key pathway *MS*0 → *MS*3 → *MS*4 earlier reported by Fabritiis and coworkers. ^7^

The free energy of binding 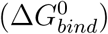, as estimated from the MSM-derived stationary populations of bound and unbound macro states of trypsin/benzamidine system -5.65 kcal/mol (with the 95% confidence interval of [-5.38,-6.02]kcal/mol) comes in reasonable agreement with experimental measurement (−6.20 kcal/mol). ^7,41^ The MSM-derived on-rate constant (3.68× 10^8^ *M* ^−1^*s*^−1^) also shows reasonable agreement with experimental report (0.29× 10^8^ *M* ^−1^*s*^−1^).^7,33^ These semi-quantitative consistencies between simulated and experimentally measured macroscopic binding constants reflect fidelities of coarse-grained simulation in aptly describing biomolecular recognition process.

The successful venture of exploring unbiased bio-molecular recognition processes in trypsin-benzamidine system using coarse-grained model encouraged us to assess the ability of our strategy in hierarchically more complex protein/ligand system namely L99A T4 Lysozyme/benzene. This system has been the subject of recent computational investigations, albeit in atomistic representations, where multiple pathways of ligand access to the cavity of L99A T4 Lysozyme have been observed.^8,36,42,43^ We wanted to explore if the coarse-grained model of this system can recover the multiple pathways. While optimizing the elastic network for L99A T4 Lysozyme, we observed that application of homogeneous network has different effect on different domain of the protein. We found that C-terminal domain (Helix4 to Helix9) was becoming more rigid (region where the cavity resides) while N-terminal domain (Helix1 to Helix3) was more flexible (This is true in atomistic simulation as well). Therefore, we focused on the optimisation the elastic network of C-terminal domain of L99A T4 Lysozyme so as to regain its flexibility and maintain its tertiary fold of all the protein. Accordingly we compared the backbone RMSD for C-terminal domain (Helix4 to Helix9) of atomistic and coarse grain simulations (see figure S5 in SI) and simulated the diffusion of benzene in an aqueous solution of L99A T4 Lysozyme within coarse-grained frame-work. We found that in our coarse-grained model, benzene exhaustively explores the solvent media and protein surface and eventually finds and spontaneously binds the solvent-occluded cavity of L99A T4 Lysozyme. (See figure S6 in SI for binding profile and figure 4A) for an overlay of simulated and crystallographic binding pose).

**Figure 4:**
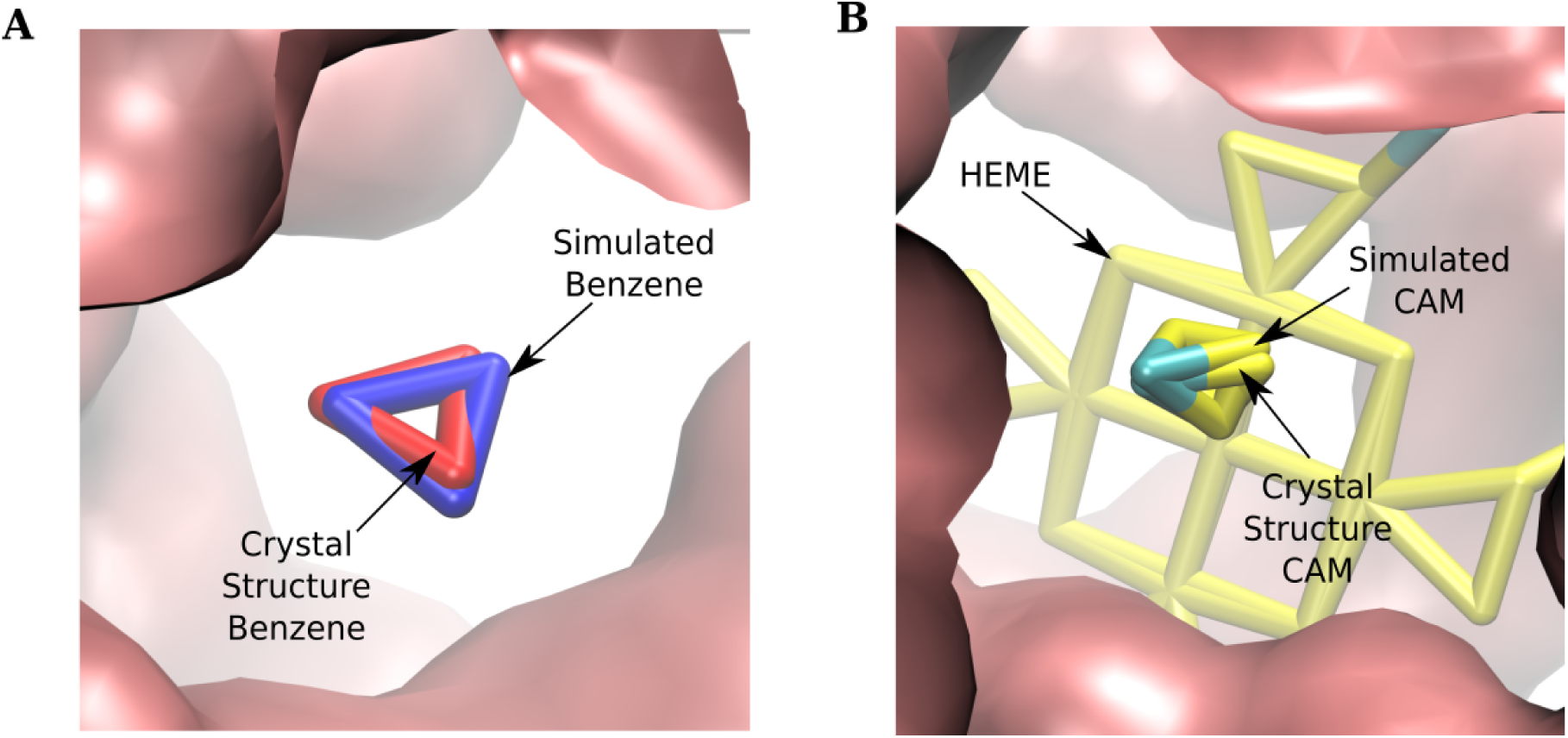
Close view of ligand pose after superimposing backbones of crystal structure and simulated structure (A) for L99A T4lysozyme Benezne system (B) for Cytochrome p450cam system.

The affordable computational cost allowed us to perform a large set of multi-microsecond long binding trajectories involving L99A T4 Lysozyme and benzene. The inspection of these trajectories revealed multiple pathways of benzene binding to the cavity of L99A T4 Lysozyme (see figure 5). Movie S3 shows the glimpses of all five pathways observed in the coarse-grained simulations. Specifically, our coarse-grained simulation trajectories revealed five binding pathways: three of the binding pathways (involving helix4-6,helix5-6-7 and helix7-helix9 of L99A T4 Lysozyme) (figure 5 A-C) are reminiscent of binding pathways observed in unbiased all-atom simulation trajectories by Mondal et al. ^8^ On the other hand, a subset of coarse-grained trajectories also re-discover two additional binding pathways, involving helix3-5 and helix4-5 (figure 5 D-E), previously hypothesised in biased simulations.^36^ While the observations of spontaneous recognition in coarse-grained simulations of trypsin-benzamidine and L99A T4 Lysozyme/benzene are encouraging, we stress that a ready-made protocol based upon *automated* optimisation of elastic network parameter might not work for all protein-ligand system and local supervision is needed. This is illustrated in the classic case of cytochrome P450/camphor recognition process. ^34^ We found a brute-force implementation of elastic network model does not reproduce flexibility observed in all-atom models. Subsequent close-view inspection of substrate-free protein in its elastic network MARTINI model, pointed towards a reduction in the cavity volume above the I helix and the heme because of slight mis-positioning of heme. This was also reflected by the lack of binding even after reaching metastable state for long time coarse-grained simulations. We resolved this issue by introducing additional elastic network around the heme active site, which ensured close agreement in RMSD between all-atom ^9^ and coarse-grained cytochrome P450 (see Figure 6A-B).

**Figure 5:**
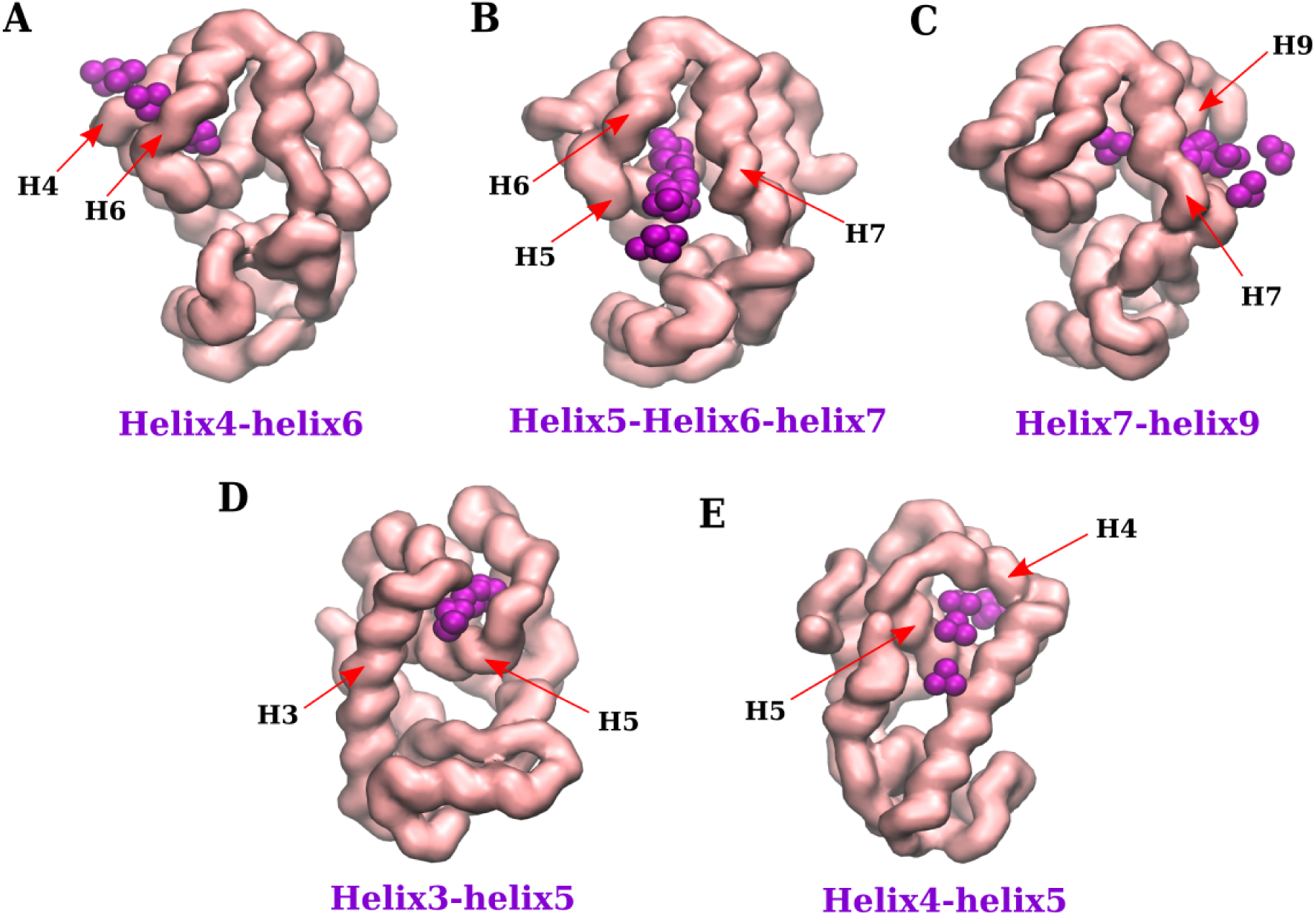
The benzene binding pathways in T4Lysozyme (A) Through Helix4-Helix6. (B) Through Helix5-Helix6-Helix7. (C) Through Helix7-Helix9 binding. (D)Through Helix3-Helix5. (E) Through Helix4-Helix5. Also see Movie S3.

**Figure 6:**
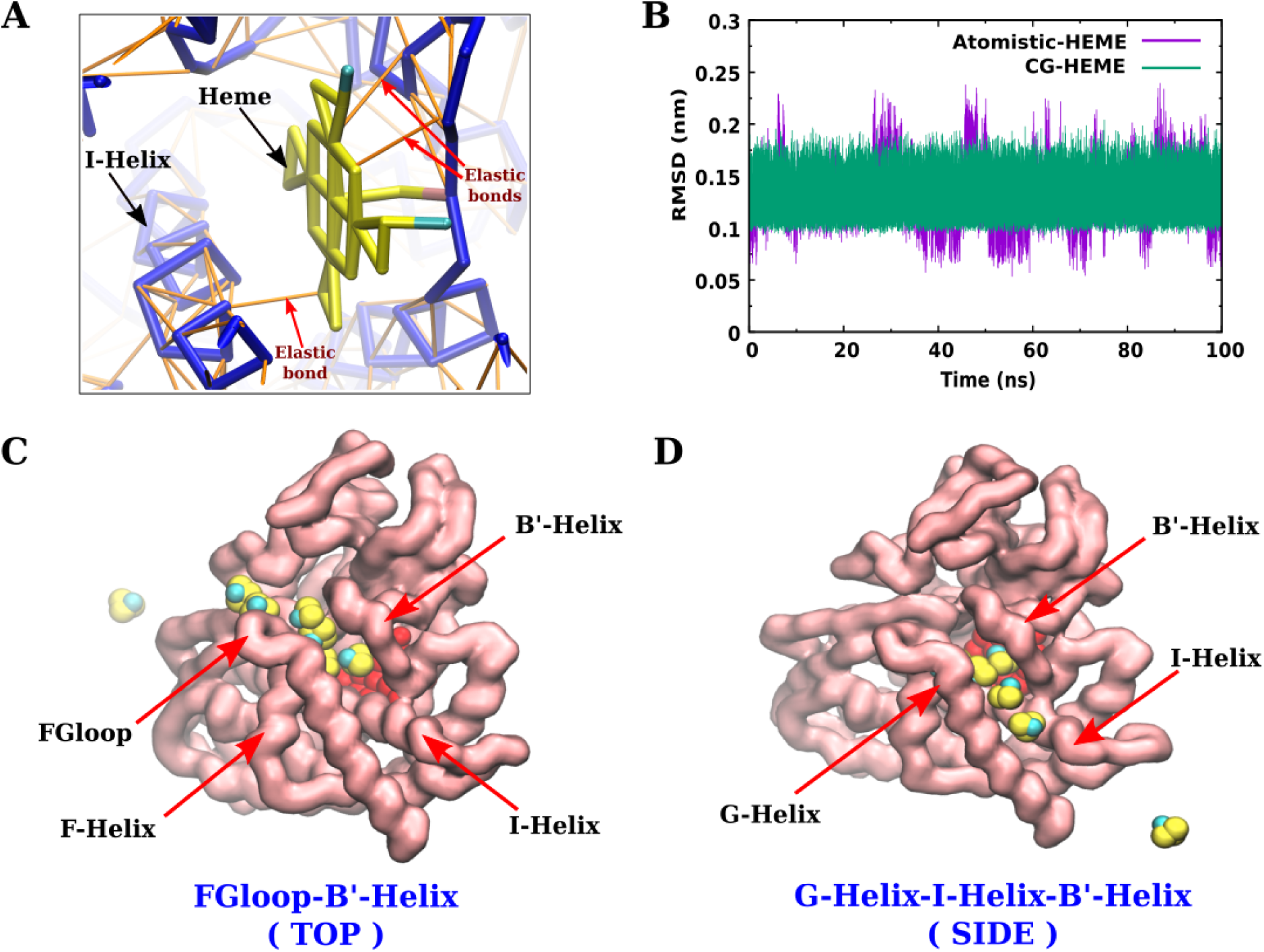
(A) Repositioning of heme using elastic network. (B) Mapping of CG heme with Atomistic. Camphor Binding Pathways (C) through FGloop and B’-Helix “TOP” (D) through G-Helix-I-Helix and B’-Helix “SIDE”. Also see Movie S4.

We found that the coarse-grained cytochrome P450, thereby optimized with locally supervised additional elastic networks, is able to recognize the camphor as a substrate efficiently. As demonstrated in movie S4, camphor eventually finds the crevices between F/G helix of cytochrome P450 in the coarse-grained trajectory and successfully lands on the heme-containing active site. Figure 4B provides the overlay of crystallographic and final simulated snapshot of cytochrome P450/camphor and figure S7 depicts the binding time profile. The short wall-clock time of the coarse-grained simulation allowed us to run a large set of independent binding simulation trajectories to explore the diversities in binding pathways. Movie S4 depicts both the camphor-P450 recognition pathways observed in coarse grained simulations. Quite gratifyingly, the major substrate-recognition pathway observed in these coarse-grained simulations via crevices of F/G loop and B’-Helix (mentioned as a “TOP” pathway in the figure 6C) is identical to what was recently observed in all-atom binding simulations of Cytochrome P450/camphor^9^, albeit at orders of magnitude larger computational cost. Moreover, a subset of these coarse-grained simulations also provided insights into other possible pathway. (“SIDE” pathway in figure 6D). More surprisingly both these pathways are already shown to exist based on the simulation studies of available crystal structures. ^47–49^ This pathway shows the entry of camphor through the crevices of G/I Helix and B’-Helix. Again more importantly we see that the simple coarse grained representation of camphor is able to locate the binding pose and the pathways reported earlier in literatures.

## Conclusion

In summary, we have demonstrated that coarse-grained simulation with optimised elastic network is a practical way forward for capturing the protein-ligand recognition pathways within affordable computational expense. Via systematic benchmarking of the coarse-grained model against atomistically simulated ligand-free protein’s tertiary fold, we have shown that coarse-grained models can potentially live upto the expectation of deciphering underlying mechanism of protein-ligand binding processes. Table 1 provides a quantitative testimony of the simulations that we could afford using coarse-grained model. The number of direct binding trajectories that these coarse-grained models enabled us to explore, from solvent to cavity, is unprecedented in all-atom simulations. The significantly lower computational cost associated with coarse-grained simulation also provides opportunities for better statistics and potential discovery of new, therapeutically relevant recognition pathways. Most importantly, the kinetic on-rate constants as estimated from these coarse-grained simulations comes within reasonable agreement with that of experiment, which eventually provides leverage for exploring metabolite and small molecule recognitions in large-scale biological systems.

This approach being computationally inexpensive, provides a practical way-forward for replacing traditional docking based small molecule discovery by high-throughput coarse-grained simulation for potential binding site and inhibitor discovery. The possibility of exhaustive ligand sampling around protein in coarse-grained model allows one to explore statistically meaningful ligand-binding hotspots, thereby exploring potential allosteric sites, apart from the native binding sites. Towards this end, mixed resolution models^50,51^ with protein sites proximal to binding regions modelled with all-atom descriptions and the rest of the system treated by coarse-grained representation holds exciting promises. This opens promising windows for exploring mechanistic pathways of protein-protein association processes using coarse-grained simulation approach. Recent progresses in back mapping from multi resolution coarse-grained models^52^ also provide an option for post-facto reconstruction of the atomistically detailed mechanisms of biomolecular recognition processes.

## Materials and methods

### Atomistic simulations of apo proteins and ligands

Unbiased all-atom MD simulations of ligand-free proteins for all three systems were performed. These atomistic trajectories served as the benchmarks for the optimisation of elastic networks of the coarse-grained models. As shown in figure 1, we simulated three apo-proteins (namely, Trypsin (PDB id: 1S0Q), L99A T4Lysozyme (PDB id: 3DMV) and cytochrome P450(PDB id: 1PHC)) in their all-atom model. All simulation started with the crystal structure of protein and the simulation box was solvated with water with the protein at the centre. All protein atoms were modelled using charmm36 force field^53^ and water was modelled as TIP3P.^54^ Each individual system was simulated for 100 ns using leapfrog integrator with a time step of 2 femtosecond. All MD simulations were performed with the Gromacs 2018 simulation package^55^, in most cases benefiting from usage of Graphics processing units (GPU).^56^

Similarly, we also performed the all-atom simulations of all the ligands to obtain the benchmark parameters for optimization of coarse-grain ligand models. We obtained the ligands co-ordinated from the bound complex structures of respective systems (namely, Trypsin-benzamidine (PDB id: 3ATL), L99A T4Lysozyme-benzene (PDB id: 3DMX) and cytochrome P450-camphor (PDB id: 2CPP) and also heme cofactor). The all-atom camphor parameters were obtained from our earlier work^9^ and heme were modelled using charmm36 force field.^53^ The benzene and benzamidine molecules were modelled using general AMBER force field.^57^ The details of the system and simulation protocols are provided in SI text and table S1.

### Coarse grained unbiased binding simulation

We first mapped atomistic protein structure to the coarse-grained model using martinize.py script. Subsequently, we employed elastic networks ^32^ among all backbone atoms of coarse-grained protein^30,31^ with elastic network parameters optimised as described in main text. The ligand molecules were coarse-grained and parameterised where necessary (see SI methods). All binding simulations were initiated by placing the apo form of coarse-grained protein at the centre of a cubic box with the empty space filled with Martini water and Martini ions,^58^ keeping the sodium chloride concentration at 150 mM and rendering the system charge neutral. Multiple copies of coarse-grained ligand molecules were placed in random positions in the solvent. All ligand molecules were allowed to diffuse freely and no artificial bias was introduced throughout the simulation. All MD simulations were performed with the Gromacs 2018 simulation package,^55^ together with GPU accelerations and reaction fields, as per Martini recommendations.^59^ For an exhaustive quantification of binding pathways and computation of macroscropic binding constants, we undertook the task of building a Markov state model (MSM)^38,39^ for trypsin-benzamidine system The details of the coarse-grained system and simulation protocols are provided in SI text and table S2.

## Supporting information

Coarse-grained topologies of ligand

## Supplemental Material

Supplemental information on details of method and materials, supplemental figures, (PDF) supporting movies (mp4) and coarse-grained ligand topologies (zip)

## Acknowledgements

This work was supported by computing resources obtained from shared facility of TIFR Center for Interdisciplinary Sciences, India. JM would like to acknowledge research intramural research grants obtained from TIFR, DAE, India, Ramanujan Fellowship and Early Career Research funds provided by the Department of Science and Technology (DST) of India (ECR/2016/000672).

## References

1. Pang, X.; Zhou, H.-X. Rate Constants and Mechanisms of Protein-Ligand Binding. Annual Review of Biophysics 2017, 46, 105–130, PMID: 28375732.

2. Pan, A. C.; Borhani, D. W.; Dror, R. O.; Shaw, D. E. Molecular determinants of drug–receptor binding kinetics. Drug Discov Today 2013, 18, 667–673.

3. Shan, Y.; Kim, E. T.; Eastwood, M. P.; Dror, R. O.; Seeliger, M. A.; Shaw, D. E. How Does a Drug Molecule Find Its Target Binding Site? Journal of the American Chemical Society 2011, 133, 9181–9183, PMID: 21545110.

4. Dror, R. O.; Pan, A. C.; Arlow, D. H.; Borhani, D. W.; Maragakis, P.; Shan, Y.; Xu, H.; Shaw, D. E. Pathway and mechanism of drug binding to G-protein-coupled receptors. Proceedings of the National Academy of Sciences 2011, 108, 13118–13123.

5. Kmiecik, S.; Gront, D.; Kolinski, M.; Wieteska, L.; Dawid, A. E.; Kolinski, A. Coarse-Grained Protein Models and Their Applications. Chemical Reviews 2016, 116, 7898–7936, PMID: 27333362.

6. Marrink, S. J.; Tieleman, D. P. Perspective on the Martini model. Chem. Soc. Rev. 2013, 42, 6801–6822.

7. Buch, I.; Giorgino, T.; De Fabritiis, G. Complete reconstruction of an enzyme-inhibitor binding process by molecular dynamics simulations. Proc Natl Acad Sci 2011, 108, 10184–10189.

8. Mondal, J.; Ahalawat, N.; Pandit, S.; Kay, L. E.; Vallurupalli, P. Atomic resolution mechanism of ligand binding to a solvent inaccessible cavity in T4 lysozyme. PLOS Computational Biology 2018, 14, 1–20.

9. Ahalawat, N.; Mondal, J. Mapping the Substrate Recognition Pathway in Cytochrome P450. Journal of the American Chemical Society 2018, 140, 17743–17752.

10. Plattner, N.; Noe, F. Protein conformational plasticity and complex ligand-binding kinetics explored by atomistic simulations and Markov models. Nat Commun 2015, 6, 7653.

11. Dror, R. O.; Dirks, R. M.; Grossman, J. P.; Xu, H.; Shaw, D. E. Biomolecular Simulation: A Computational Microscope for Molecular Biology. Annu Rev Biophys 2012, 41, 429–452.

12. Paul, F.; Wehmeyer, C.; Abualrous, E. T.; Wu, H.; Crabtree, M. D.; Schöneberg, J.; Clarke, J.; Freund, C.; Weikl, T. R.; Noé, F. Protein-peptide association kinetics beyond the seconds timescale from atomistic simulations. Nature Communications 2017, 8.

13. Ingolfsson, H. I.; Melo, M. N.; van Eerden, F. J.; Arnarez, C.; Lopez, C. A.; Wassenaar, T. A.; Periole, X.; de Vries, A. H.; Tieleman, D. P.; Marrink, S. J. Lipid Organization of the Plasma Membrane. Journal of the American Chemical Society 2014, 136, 14554–14559, PMID: 25229711.

14. Risselada, H. J.; Marrink, S. J. The molecular face of lipid rafts in model membranes. Proceedings of the National Academy of Sciences 2008, 105, 17367–17372.

15. Bennett, W. D.; Tieleman, D. P. Water Defect and Pore Formation in Atomistic and Coarse-Grained Lipid Membranes: Pushing the Limits of Coarse Graining. Journal of Chemical Theory and Computation 2011, 7, 2981–2988, PMID: 26605486.

16. Kirsch, S. A.; Bockmann, R. A. Membrane pore formation in atomistic and coarsegrained simulations. Biochimica et Biophysica Acta (BBA) - Biomembranes 2016, 1858, 2266–2277, Biosimulations of lipid membranes coupled to experiments.

17. Periole, X.; Zeppelin, T.; Schiott, B. Dimer Interface of the Human Serotonin Transporter and Effect of the Membrane Composition. Sci. Rep. 2017, 8, 5080.

18. Yoo, J.; Cui, Q. Membrane-Mediated Protein-Protein Interactions and Connection to Elastic Models: A Coarse-Grained Simulation Analysis of Gramicidin A Association. Biophysical Journal 2013, 104, 128–138.

19. Marrink, S. J.; Mark, A. E. Molecular Dynamics Simulation of the Formation, Structure, and Dynamics of Small Phospholipid Vesicles. Journal of the American Chemical Society 2003, 125, 15233–15242, PMID: 14653758.

20. Mondal, J.; Sung, B. J.; Yethiraj, A. Sequence-Directed Organization of beta-Peptides in Self-Assembled Monolayers. The Journal of Physical Chemistry B 2009, 113, 9379–9385, PMID: 19545127.

21. Mondal, J.; Sung, B. J.; Yethiraj, A. Sequence dependent self-assembly of beta-peptides: Insights from a coarse-grained model. The Journal of Chemical Physics 2010, 132, 065103.

22. Wu, Z.; Cui, Q.; Yethiraj, A. Driving Force for the Association of Hydrophobic Peptides: The Importance of Electrostatic Interactions in Coarse-Grained Water Models. The Journal of Physical Chemistry Letters 2011, 2, 1794–1798.

23. Baoukina, S.; Tieleman, D. P. Direct Simulation of Protein-Mediated Vesicle Fusion: Lung Surfactant Protein B. Biophysical Journal 2010, 99, 2134–2142.

24. Markvoort, A. J.; Marrink, S. J. In *Chapter 11 - Lipid Acrobatics in the Membrane Fusion Arena*; Chernomordik, L. V., Kozlov, M. M., Eds.; Current Topics in Membranes; Academic Press, 2011; Vol. 68; pp 259–294.

25. Levi, M.; Noel, J. K.; Whitford, P. C. Studying ribosome dynamics with simplified models. Methods 2019, 162-163, 128–140, Experimental and Computational Techniques for Studying Structural Dynamics and Function of RNA.

26. Shimizu, M.; Takada, S. Reconstruction of Atomistic Structures from Coarse-Grained Models for Protein-DNA Complexes. Journal of Chemical Theory and Computation 2018, 14, 1682–1694, PMID: 29397721.

27. Brandani, G. B.; Niina, T.; Tan, C.; Takada, S. DNA sliding in nucleosomes via twist defect propagation revealed by molecular simulations. Nucleic Acids Research 2018, 46, 2788–2801.

28. Brandner, A.; Schaller, A.; Melo, F.; Pantano, S. Exploring DNA dynamics within oligonucleosomes with coarse-grained simulations: SIRAH force field extension for protein-DNA complexes. Biochemical and Biophysical Research Communications 2018, 498, 319–326, Multiscale Modeling.

29. Marrink, S. J.; Risselada, H. J.; Yefimov, S.; Tieleman, D. P.; de Vries, A. H. The MARTINI Force Field:a Coarse Grained Model for Biomolecular Simulations. The Journal of Physical Chemistry B 2007, 111, 7812–7824, PMID: 17569554.

30. Monticelli, L.; Kandasamy, S. K.; Periole, X.; Larson, R. G.; Tieleman, D. P.; Marrink, S.-J. The MARTINI Coarse-Grained Force Field: Extension to Proteins. Journal of Chemical Theory and Computation 2008, 4, 819–834, PMID: 26621095.

31. de Jong, D. H.; Singh, G.; Bennett, W. F. D.; Arnarez, C.; Wassenaar, T. A.; Schafer, L. V.; Periole, X.; Tieleman, D. P.; Marrink, S. J. Improved Parameters for the Martini Coarse-Grained Protein Force Field. Journal of Chemical Theory and Computation 2013, 9, 687–697, PMID: 26589065.

32. Periole, X.; Cavalli, M.; Marrink, S.-J; Ceruso, M. A. Combining an Elastic Network With a Coarse-Grained Molecular Force Field: Structure, Dynamics, and Intermolecular Recognition. Journal of Chemical Theory and Computation 2009, 5, 2531–2543, PMID: 26616630.

33. Guillain, F.; Thusius, D. Use of proflavine as an indicator in temperature-jump studies of the binding of a competitive inhibitor to trypsin. Journal of the American Chemical Society 1970, 92, 5534–5536, PMID: 5449454.

34. Raag, R.; Poulos, T. L. Crystal structures of cytochrome p-450CAM complexed with camphane, thiocamphor, and adamantane, factors controlling P-450 substrate hydroxylation. Biochemistry 1991, 30, 2674–2684.

35. Eriksson, A. E.; Baase, W. A.; Wozniac, J. A.; Matthews, B. W. A cavity-containing mutant of T4 lysozyme is stabilized by buried benzene. Nature 1992, 355, 371–373.

36. Capelli, R.; Carloni, P.; Parrinello, M. Exhaustive Search of Ligand Binding Pathways via Volume-Based Metadynamics. The Journal of Physical Chemistry Letters 2019, 10, 3495–3499.

37. Yamane, J.; Yao, M.; Zhou, Y.; Hiramatsu, Y.; Fujiwara, K.; Yamaguchi, T.; Yamaguchi, H.; Togame, H.; Tsujishita, H.; Takemoto, H.; Tanaka, I. In-crystal affinity ranking of fragment hit compounds reveals a relationship with their inhibitory activities. Journal of Applied Crystallography 2011, 44, 798–804.

38. Bowman, G. R.; Pande, V. S.; Noé, F. An Introduction to Markov State Models and Their Application to Long Timescale Molecular Simulation. Advances in Experimental Medicine and Biology. 2014.

39. Chodera, J. D.; Noé, F. Markov state models of biomolecular conformational dynamics. Current Opinion in Structural Biology 2014, 25, 135–144.

40. Scherer, M. K.; Trendelkamp-Schroer, B.; Paul, F.; Perez-Hernandez, G.; Hoffmann, M.; Plattner, N.; Wehmeyer, C.; Prinz, J.-H; Noe, F. PyEMMA 2: A Software Package for Estimation, Validation, and Analysis of Markov Models. Journal of Chemical Theory and Computation 2015, 11, 5525–5542.

41. Mares-guiai, M.; Shaw, E. Studies On the Active Center of Trypsin. The Binding of Amidines And Guanidines As Models of the substrate side chain. J.Biol.Chem. 1965, 240.

42. Lamim Ribeiro, J. M.; Tiwary, P. Toward Achieving Efficient and Accurate Ligand-Protein Unbinding with Deep Learning and Molecular Dynamics through RAVE. Journal of Chemical Theory and Computation 2019, 15, 708–719.

43. Ariane N., D., Zuckerman; Arantes, G. M. Escape of a Small Molecule from Inside T4 Lysozyme by Multiple Pathways. Biophysical J. 2018, 114, 1058–1066.

44. Feher, V. A.; Baldwin, E. P.; Dahlquist, F. W. Access of ligands to cavities within the core of a protein is rapid. Nat. Struct. Mol. Biol 1996, 3, 516–521.

45. Griffinf, B. W.; Peterson, J. A. Camphor Binding by. 1972, 11, 4740–4746.

46. de Montellano, P. R. O. Cytochrome P450: Structure, Mechanism, and Biochemistry.

47. Lademann, S. K.; Lounnas, V.; Wade, R. C. How do substrates enter and products exit the buried active site of cytochrome P450cam? 2. Steered molecular dynamics and adiabatic mapping of substrate pathways11Edited by J. Thornton. Journal of Molecular Biology 2000, 303, 813–830.

48. Cojocaru, V.; Winn, P. J.; Wade, R. C. The ins and outs of cytochrome P450s. Biochimica et Biophysica Acta (BBA) - General Subjects 2007, 1770, 390–401, P450.

49. Urban, P.; Lautier, T.; Pompon, D.; Truan, G. Ligand Access Channels in Cytochrome P450 Enzymes: A Review. International Journal of Molecular Sciences 2018, 19.

50. Wassenaar, T. A.; Ingolfsson, H. I.; Priea, M.; Marrink, S. J.; Schafer, L. V. Mixing Martini: Electrostatic Coupling in Hybrid atomistic and coarse-grained biomolecular simulation. The Journal of Physical Chemistry B 2013, 117, 3516–3530.

51. Zavadlav, J.; Melo, M. N.; Marrink, S. J.; Praprotnik, M. Adaptive resolution simulation of an atomistic protein in MARTINI water. The Journal of Chemical Physics 2014, 140, 054114.

52. Peng, J.; Yuan, C.; Ma, R.; Zhang, Z. Backmapping from Multiresolution Coarse-Grained Models to Atomic Structures of Large Biomolecules by Restrained Molecular Dynamics Simulations Using Bayesian Inference. Journal of Chemical Theory and Computation 2019, 15, 3344–3353, PMID: 30908042.

53. Best, R. B.; Zhu, X.; Shim, J.; Lopes, P. E. M.; Mittal, J.; Feig, M.; MacKerell, A. D. Optimization of the Additive CHARMM All-Atom Protein Force Field Targeting Improved Sampling of the Backbone ?, *ψ* and Side-Chain *χ*^1^ and *χ*^2^ Dihedral Angles. J.Chem.Theory Comput. 2012, 8, 3257–3273.

54. Jorgensen, W. L.; Chandrasekhar, J.; Madura, J. D.; Impey, R. W.; Klein, M. L. Comparison of simple potential functions for simulating liquid water. The Journal of Chemical Physics 1983, 79, 926–935.

55. Abraham, M. J.; Murtola, T.; Schulz, R.; Pall, S.; Smith, J. C.; Hess, B.; Lindahl, E. GROMACS: High performance molecular simulations through multi-level parallelism from laptops to supercomputers. SoftwareX 2015, 1-2, 19–25.

56. Kutzner, C.; Pall, S.; Fechner, M.; Esztermann, A.; de Groot, B. L.; Grubmaeller, H. Best bang for your buck: GPU nodes for GROMACS biomolecular simulations. J.Comput.Chem. 2015, 36, 1990–2008.

57. Wang, J.; Wolf, R. M.; Caldwell, J. W.; Kollman, P. A.; Case, D. A. Development and testing of a general amber force field. Journal of Computational Chemistry 2004, 25, 1157–1174.

58. Marrink, S. J.; de Vries, A. H.; Mark, A. E. Coarse Grained Model for Semiquantitative Lipid Simulations. The Journal of Physical Chemistry B 2004, 108, 750–760.

59. de Jong, D. H.; Baoukina, S.; Ingolfsson, H. I.; Marrink, S. J. Martini straight: Boosting performance using a shorter cutoff and GPUs. Computer Physics Communications 2016, 199, 1–7.

