## Supplemental information for "Capturing Protein-Ligand Recognition Pathways in Coarse-grained Simulation"

### Supporting information for "Capturing protein-ligand recognition pathways in coarse-grained simulation"

### Methodology

### Simulation details

#### Atomistic simulations of apo-proteins and ligands

The benchmark unbiased atomistic simulations were performed using the ligand-free form of the protein. As discussed in the main text, we have explored three protein systems, namely, Trypsin (PDB code: 1S0Q), L99A T4Lysozyme (PDB: 3DMV) and cytochrome P450(PDB: 1PHC).[1–3] All simulations started with the apo-protein in their crystallographic pose. Charmm36 force fields[4] were employed to model the protein in their all-atom representations. In each cases, the simulation box was solvated with water ( parameterized using TIP3P[5] model ), with the protein centred at the box. Sufficient numbers of sodium and chloride ions were added to keep the sodium chloride concentration at 150 mM and render the system charge neutral. Table S1 details the system sizes for all three protein simulated in the work.

Similarly, we performed benchmarked unbiased atomistic simulations of all the ligands to obtain the bond, angle and dihedral parameter distributions and compare with their coarse-grain models (see Fig S1 D). The ligand coordinates were collected from the bound crystal PDB structures of respective protein-ligand complexes, namely, Trypsin-benzamidine (PDB code: 3ATL), L99A T4Lysozyme-benzene (PDB: 3DMX) and cytochrome P450-camphor (PDB: 2CPP). We also simulated the heme group, which is the cofactor of CYP450 protein. The all-atom camphor parameters were obtained from our earlier work[6]and heme were modelled using charmm36 force field.[4] The benzene and benzamidine molecules were modelled using general AMBER force field[7] with the help of ACPYPE[8] topology generator script.

All simulations were performed using NPT ensemble. The average temperature being maintained at 303 K using Nose Hoover thermostat[9,10] with a relaxation time of 1 ps. The average isotropic pressure of 1 bar was maintained using Parrinello-Rahman barostat.[11]

The Verlet cutoff scheme[12] was employed throughout the simulation with the Lenard Jones interaction extending to 1.2 nm and long-range electrostatic interactions treated by Particle Mesh Ewald summation.[13] All bond lengths involving hydrogen atoms of the proteins and the ligands were constrained using the LINCS algorithm and water molecules were kept rigid using the SETTLE[14] approach. Simulations were performed for 100 ns using the leapfrog integrator with a time step of 2 fs and initiated by randomly assigning the velocities of all particles.

#### Coarse-grained parameterisation of ligand

The coarse grained topology and parameters for all ligands were obtained by adopting protocols provided by martini forcefield homepage (http://www.cgmartini.nl/index.php/tutorials-general-introduction/parametrzining-new-molecule) using both approaches 1. Parameterizing a new molecule based on known fragments and 2. Parametrizing a new molecule based on atomistic simulations. For benzamidine we used first approach of parameterizations based on known fragments. The mapping scheme is shown in Figure S1 A-C. The bond, angle and dihedral parametrization were performed via iterative comparison with all atom counterparts (see Fig S1 D). For coarse-grained benzene, we used same topology and parameters as that of coarse-grain benzene solvent[15] and reparameterized the bond lengths based on atomistic simulation. For camphor we used second approach of parametrizing a new molecule based on atomistic simulations. Here we made use of auto-martini[16] to get an idea of mapping and bead type and to generate coarse-grain model of camphor. We then tested this auto-generated camphor topology but in the simulation we observed that the this camphor topology was failing to maintain spherical shape similar to atomistic camphor. Therefore, we then modified this auto-generated topology based on our own atomistic mapping keeping same bead types. For coarse-grained heme we used same cofactor topology provided by Marrink et. al.[17] and slightly tuned the bond lengths based on our mapping from atomistic heme simulation. All ligand topologies and itp files are provided with the communication.

#### Coarse grained binding simulation

In all three systems of our interest, the X-ray crystallographic structures of ligand-free protein were used as the starting points for all of the binding simulations. We first mapped atomistic protein structure to the coarse-grained model using martinize.py script. Subsequently, we employed elastic networks[18] among all backbone atoms of coarse-grained protein[15,19] with elastic network parameters optimised as described in main text. All binding simulations were initiated by placing the apo form of coarse-grained protein at the centre of a cubic box with the empty space filled with Martini water and Martini ions,[20] keeping the sodium chloride concentration at 150 mM and rendering the system charge neutral. Multiple copies of coarse-grained ligand molecules were placed in random positions in the solvent. All ligand molecules were allowed to diffuse freely and no artificial bias was introduced throughout the simulation. Table S2 provides the details of coarse-grained system sizes for all three systems of interest.

All MD simulations were performed with the Gromacs 2018 simulation package,[21] in most cases benefiting from usage of Graphics processing units.[22] We first energy minimized the systems using steepest descent algorithm to remove any bad contacts. We first performed short 500 ns long equilibration with 2 fs timestep in NVT ensemble, with the average temperature being maintained at 303K via berendsen thermostat.[23] Subsequently, we performed multiple stages of equilibration, each 500 ns long, with time steps systematically increasing from 2 fs , 10 fs and finally to 20 fs in NPT ensemble, with the average temperature maintained at 303K using the Velocity-rescale thermostat[24] with a relaxation time of 1.0 ps and an average isotropic pressure of 1 bar maintained with the Berendsen barostat.[23] The Verlet cutoff[12] scheme was employed throughout the simulation with the Lenard Jones interaction extending to 1.1 nm and long-range electrostatic interactions treated by reaction-field method as per recommendations of MARTINI protocols.[25] The production runs were performed using the leapfrog integrator with a time step of 20 fs and initiated by randomly assigning the velocities of all particles. We performed multiple independent runs each of 10 $\mu$s long, differing in the assignment of initial velocity seeds. Then trajectories were investigated for binding by calculating the radial distance between respective centres of mass of ligand and protein cavity and the root-mean-squared deviation (RMSD) of the simulated conformation of the coarse-grained protein-ligand system from that of coarse-grained crystallographic bound pose. The ligand was ascertained to be bound to the cavity when the RMSD remained below 0.2 nm and the cavity-ligand distance was below 0.7 nm for a simulation duration of at least 100 ns.

#### Markov state model analysis of CG Trypsin-Benzamidine binding simulations

For an exhaustive quantification of binding pathways and computation of macroscropic binding constants, we undertook the task of building a Markov state model (MSM)[26,27] for trypsin-benzamidine system. For this purpose, apart from the 30 copies of 10 microsecond-long trajectories discussed above, we additionally performed 26 short independent trajectories, each 1 microsecond long plus 526 short independent trajectories, each 100ns in duration plus 2000 short independent trajectories, each 10 ns in duration. These short trajectories were initiated from different intermediates observed from the long binding simulations and improved the statistics of the MSM that was derived from analysis of the MD trajectories (see below). Overall, an aggregate of about 128 $\mu$s of CG unbiased trajectories was recorded for building MSM of trypsin-benzamidine system.

We employed PyEMMA[28](http://pyemma.org) to construct and analyze the MSM obtained from the combination of all the recorded MD trajectories of Trypsin benzamidine system. We fitted the protein backbone from the trajectory to the backbone of its initial input structure. Then we used xyz coordinates of the ligands of the fitted trajectories as input collective variables. We employed the k-mean clustering algorithm[29] to discretize the input data into 500 clusters. Then 500 microstates MSMs were constructed at variable lag times to identify the appropriate lag time. A lag time of 20 ns was used to construct the final MSM as the implied time scale leveled off at about 20 ns which ensures the Markovianity of the model (see Fig S4 B) in SI). The implied time scale plot also suggested presence of total 5 metastable states at 20 ns lag time. We have also validated the five state MSM at 20ns by performing Chapman-Kolmogorov test ( Figure S4 C). To clearly understand the dynamics of ligand binding a coarse-grained kinetic model with five metastable states was constructed using a hidden Markov model (HMM).[30] The kinetic parameters reported in Table 1 of main text were calculated using the mean first passage time (MFPT) obtained from the MSM analysis based on five derived macrostates. The MFPT was computed as the average time taken for the transition from the initial(unbound) to the final (bound) macrostate. The calculation included both the direct transition from the initial state to the final state and transitions through other intermediate states. The on-rate and off-rate constants are respectively calculated as $k_{on} = \frac{1}{MFPT_{on}C}$ and $k_{off} = \frac{1}{MFPT_{off}}$, where C is the benzamidine concentration, 11 mM. The binding free energies (ΔG) were calculated based on the stationary populations of bound ($\pi_{bound}$) and unbound macrostates ($\pi_{unbound}$) as obtained from the MSM. $\Delta G = -RTlog\frac{\pi_{bound}}{\pi_{unbound}}$ We subsequently converted the result to a standard free energy $\Delta G^0$ for comparison with experiment. Finally, the transition path theory as proposed by Vanden-Eijnden and coworkers[31–33] was employed to calculate the transition path fluxes among the macro states.

#### Estimation of binding on-rates for L99A T4 Lysozyme and Cytochrome P450 system

In the L99A T4 Lysozyme Benzene binding simulations we observed 17 binding events and the target was unbound for 232.13 $\mu$s out of 300$\mu$s total simulation time. The binding frequency was coming out to be 0.073$\mu s^{-1}$ and considering the benzene concentration of 8.9mM in the simulation box results in an on-rate of $8.22 \times 10^6 s^{-1}M^{-1}$. Similarly, for Cytochrome P450 we observed 6 binding events and total 270 $\mu$s of unbound target resulting in binding frequency of about 0.022 $\mu s^{-1}$. Considering camphor concentration of 26 mM results in an on-rate of $0.34 \times 10^7 s^{-1}M^{-1}$. These on-rates are reported after multiplying by 4 to the final on-rates, considering an effect of slow diffusion of heavy CG water beads. Table 2 in main text provides the rate estimate from the simulations and its comparison with experimental reports.

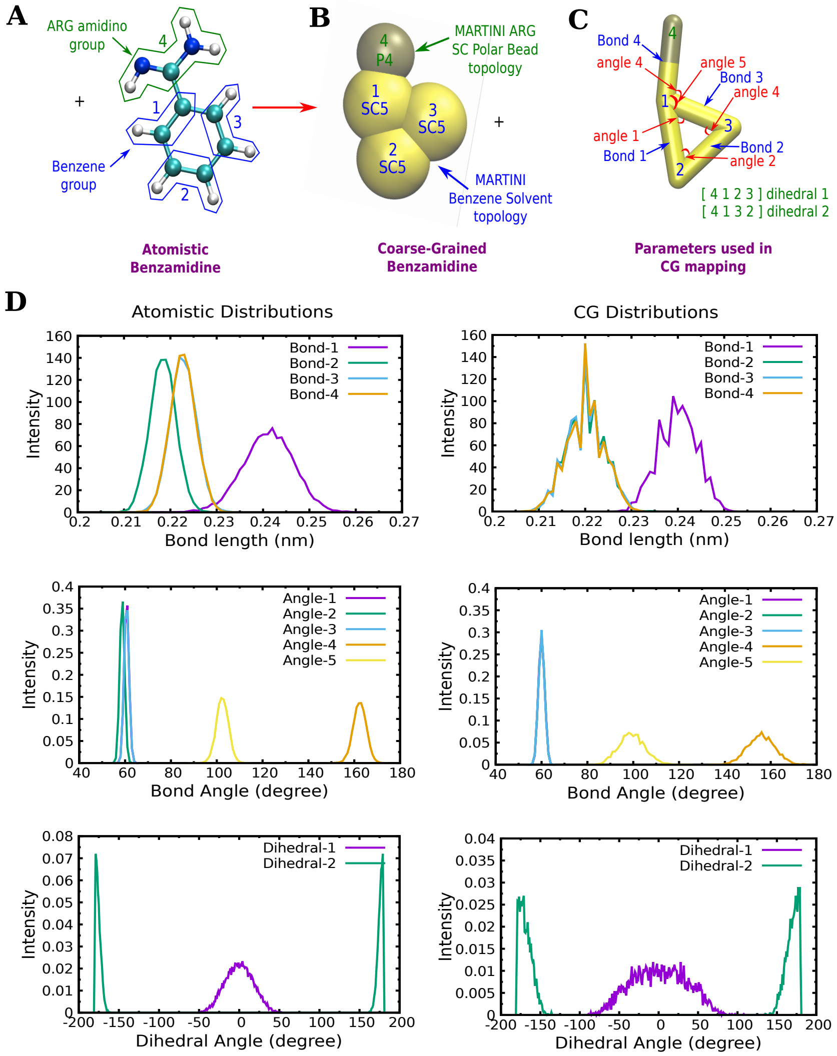


Figure S1: Parametrising a benzamidine molecule based on known fragments: (A) The atomistic benzamidine with its encircled atoms which are mapped together as a one CG bead in its corresponding coarse grained benzamidine (B). (C) The description of parameters used in mapping of Bezamidine ligand (D) The comparison of parameter distributions for CG benzamidine with the atomistic benzamidine.

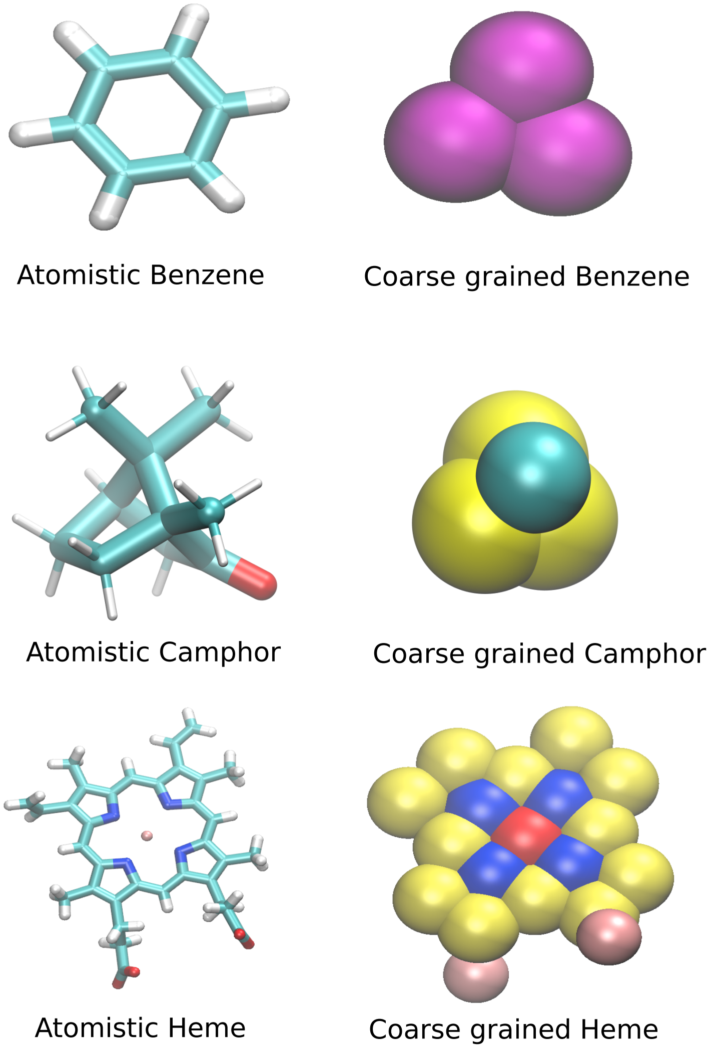


Figure S2: Other Ligands and Heme site used in the simulations in their atomistic and coarse-grained representations

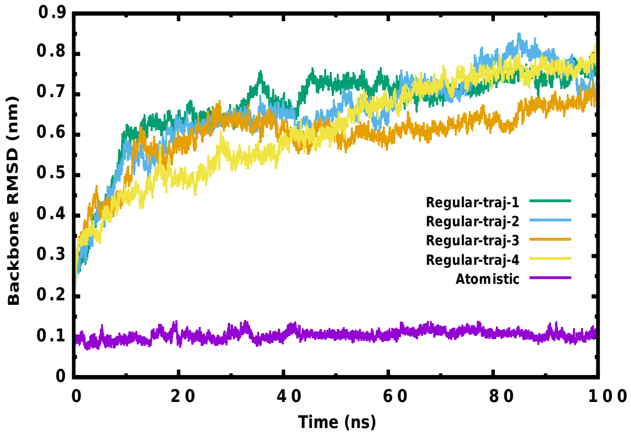


Figure S3: Backbone RMSD of CG trypsin with regular martini

Table S1: Atomistic simulation box details of apo proteins

| Protein System | PDB ID | Total No. of Particles in the Box | Cubic Box Dimesions ($nm$) |
|---|---|---|---|
| Trypsin | 1S0Q | 30432 | 6.688 |
| T4Lysozyme | 3DMV | 40896 | 7.354 |
| CYP450 | 1PHC | 70961 | 8.843 |

Table S2: Coarse grain binding simulation box details of CG protein ligand systems

| Coarse grain Protein-Ligand System | Elastic network parameter (Rc,Kf) used | Total No. of CG Particles in the box | Cubic Box Dimesions ($nm^3$) | No. of CG Ligands used | CG Ligand Concentarion in the box |
|---|---|---|---|---|---|
| Trypsin-Benzamidine | (0.90,500) | 2755 | 6.707 | 2 | 11 mM |
| T4Lysozyme-Benzene | (0.90,100) | 3314 | 7.208 | 4 | 8.9mM |
| CYP450-Camphor | (0.85,100) | 5448 | 8.842 | 8 | 26mM |

Table S3: Characteristics of the all five MSM macrostates

| State | Stationary populations % | Lifetime $\mu$s | Committor Probability | Residues Interacting with |
|---|---|---|---|---|
| MS0 | 3.05 | 0.198 | 0.0 | - |
| MS1 | 13.25 | 0.438 | 0.960 | V90,H91,W215 |
| MS2 | 6.06 | 0.222 | 0.966 | Y40,F42,K61 |
| MS3 | 30.96 | 0.946 | 0.989 | K167,Q173,K226 |
| MS4 | 44.08 | 1.578 | 1.0 | D194,Q192,S195 |

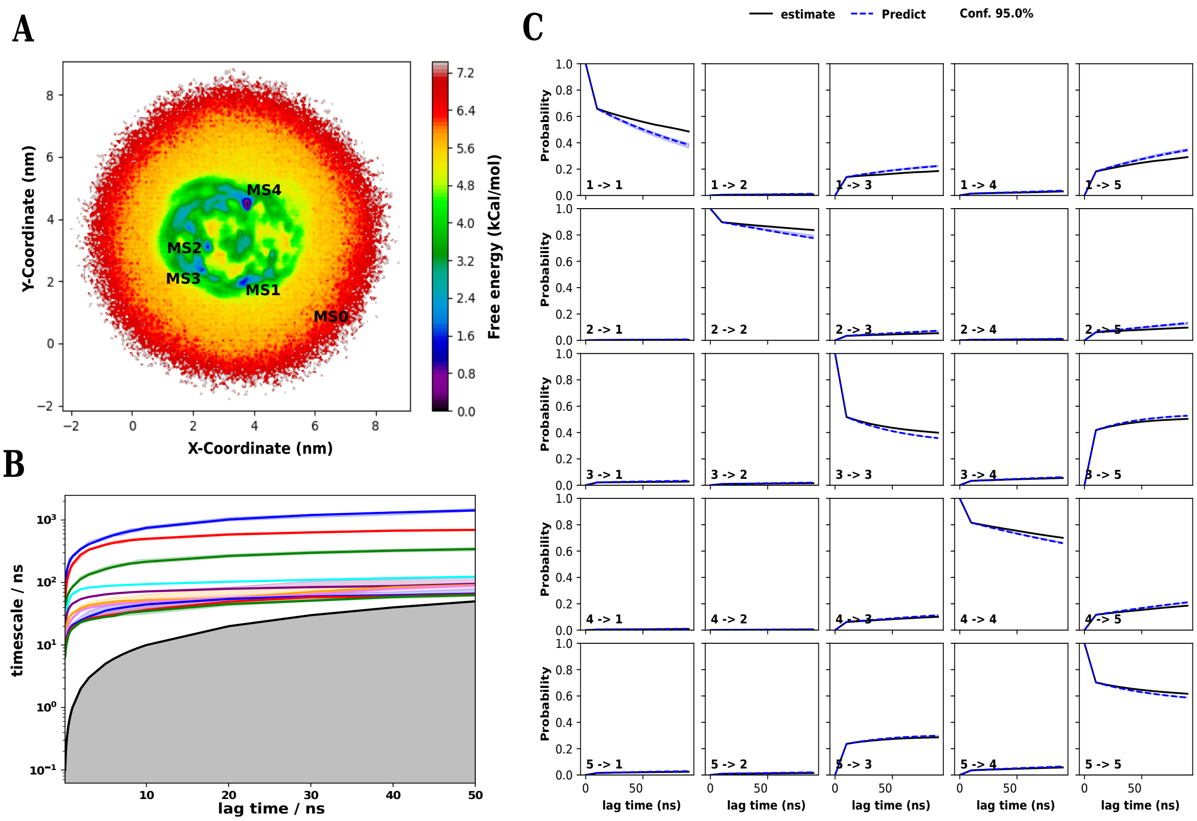


Figure S4: Development of Markov-state model for coarse-grained simulations of trypsin-benzamidine recognition process: (A) Free energy landscape along XY coordinate of benzamidine ligand shows the presence of 5 distinct minimas. The minimas are labeled based on the XY coordinate values of extrated macrostates from MSM (B) Implied timscale plot leveled off at about 20 ns and shows the presence of five distinct macrostates. (C) Chapman-Kolmogorov-test for 5 state model shows agreement between estimated and predicted MSM at the lag time of 20ns.

- Movie S1 demonstrates representative conformational change and eventual collapse of binding cavity of trypsin when using Martini forcefield without elastic network.
- Movie S2 is a representative movie of benzamidine binding to trypsin in coarse-grained model with optimised elastic network.
- Movie S3 is a movie for all five benzene binding pathways to a L99A T4lysozyme in coarse-grained model with optimised elastic network.
- Movie S4 is a movie for two different camphor binding pathways to a CytochromeP450

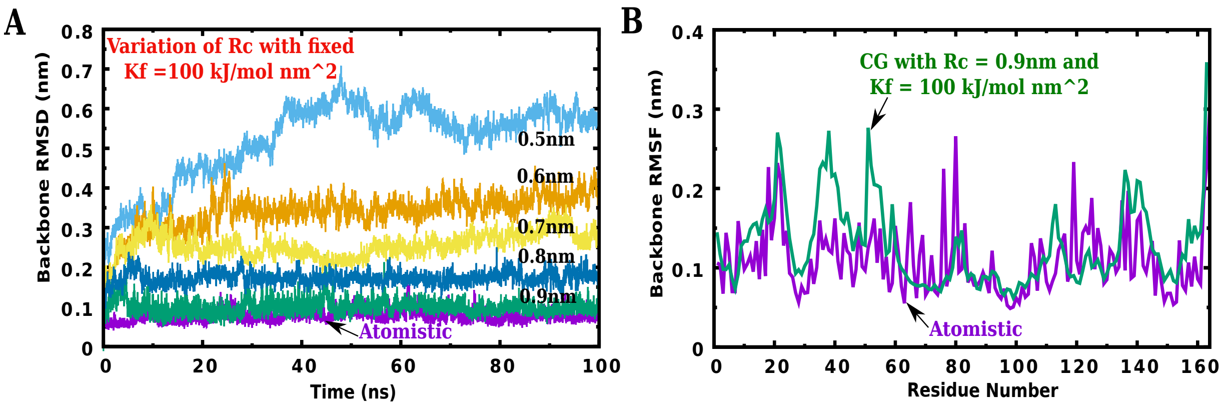


Figure S5: Optimization of elastic network for t4lysozyme and validation using RMSD C-terminal domain (Helix4 to Helix9) and RMSF of all the backbone.

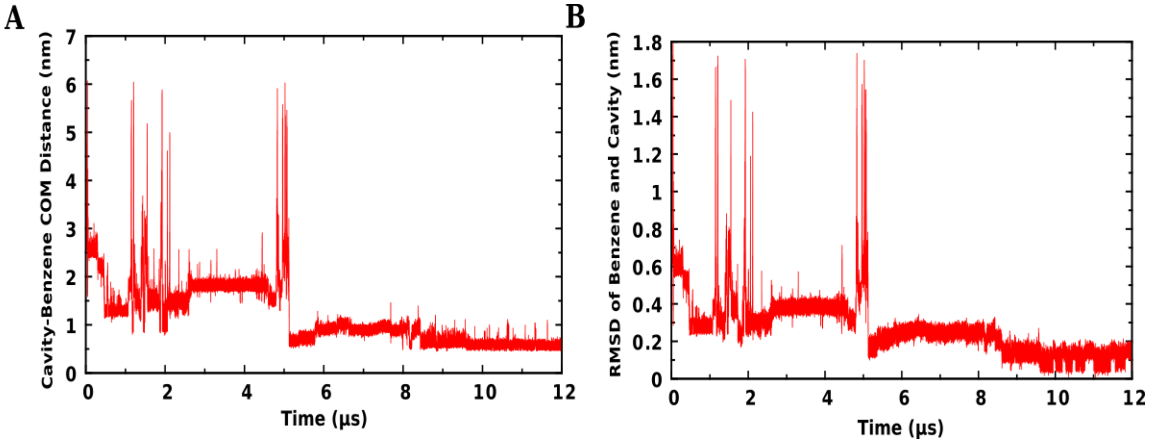


Figure S6: (A) Distance between com (center of mass) of cavity and com of Benzene (B) RMSD of benzene along with the residues within 6 $\mathring{A}$ of benzene (relative to crystallographic pose) for t4lysozyme-benzene system.

in coarse-grained model with optimised elastic network.

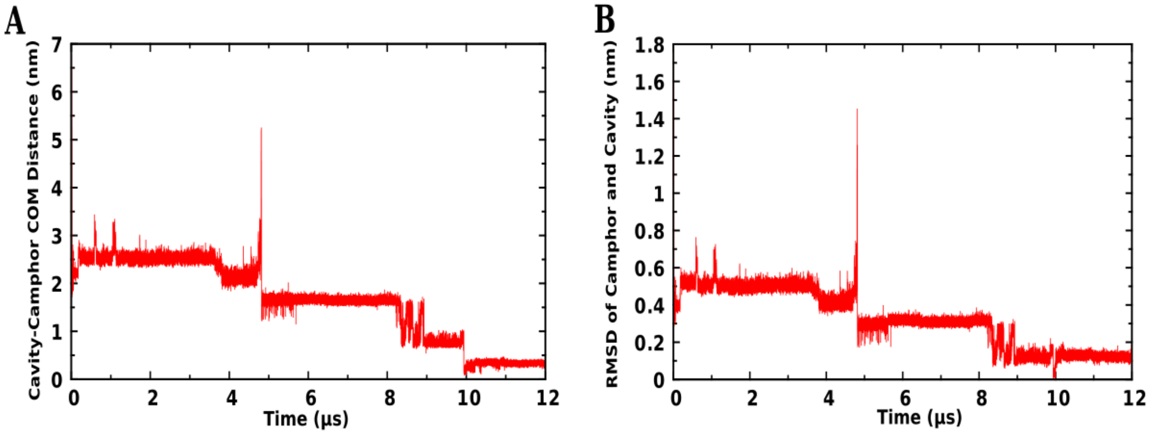


Figure S7: A) Distance between com (center of mass) of cavity and com of camphor (B) RMSD of camphor along with the residues within 6 $\mathring{A}$ of camphor (relative to crystallographic pose) for p450cam system.

### References


(1) Guillain, F.; Thusius, D. Use of proflavine as an indicator in temperature-jump studies of the binding of a competitive inhibitor to trypsin. *Journal of the American Chemical Society* **1970**, *92*, 5534–5536, PMID: 5449454.

(2) Raag, R.; Poulos, T. L. Crystal structures of cytochrome p-450CAM complexed with camphane,thiocamphor,and adamantane,factors controlling P-450 substrate hydroxylation. *Biochemistry* **1991**, *30*, 2674–2684.

(3) Eriksson, A. E.; Baase, W. A.; Wozniac, J. A.; Matthews, B. W. A cavity-containing mutant of T4 lysozyme is stabilized by buried benzene. *Nature* **1992**, *355*, 371–373.

(4) Best, R. B.; Zhu, X.; Shim, J.; Lopes, P. E. M.; Mittal, J.; Feig, M.; MacKerell, A. D. Optimization of the Additive CHARMM All-Atom Protein Force Field Targeting Improved Sampling of the Backbone ?, $\psi$ and Side-Chain $\chi1$ and $\chi2$ Dihedral Angles. *J.Chem.Theory Comput.* **2012**, *8*, 3257–3273.

(5) Jorgensen, W. L.; Chandrasekhar, J.; Madura, J. D.; Impey, R. W.; Klein, M. L.

Comparison of simple potential functions for simulating liquid water. *The Journal of Chemical Physics* **1983**, *79*, 926–935.

(6) Ahalawat, N.; Mondal, J. Mapping the Substrate Recognition Pathway in Cytochrome P450. *Journal of the American Chemical Society* **2018**, *140*, 17743–17752.

(7) Wang, J.; Wolf, R. M.; Caldwell, J. W.; Kollman, P. A.; Case, D. A. Development and testing of a general amber force field. *Journal of Computational Chemistry* **2004**, *25*, 1157–1174.

(8) Sousa Da Silva, A. W.; Vranken, W. F. ACPYPE - AnteChamber PYthon Parser interfacE. *BMC Research Notes* **2012**, *5*, 1–8.

(9) Nosé, S. A molecular dynamics method for simulations in the canonical ensemble. *Mol. Phys.* **1984**, *52*, 255.

(10) Hoover, W. Canonical dynamics: equilibrium phase-space distributions. *Phys. Rev. A* **1985**, *31*, 1695.

(11) Parrinello, M.; Rahman, A. Polymorphic transitions in single crystals: A new molecular dynamics method. *Journal of Applied Physics* **1981**, *52*, 7182–7190.

(12) Pail, S.; Hess, B. A flexible algorithm for calculating pair interactions on {SIMD} architectures. *Comput. Phys. Comm.s* **2013**, *184*, 2641 – 2650.

(13) Darden, T.; York, D.; Pederson, L. G. *J. Chem. Phys.* **1993**, *98*, 952.

(14) Miyamoto, S.; Kollman, P. Settle: An analytical version of the SHAKE and RATTLE algorithm for rigid water models. *J. Comput. Chem.* **1992**, *13*, 952–962.

(15) de Jong, D. H.; Singh, G.; Bennett, W. F. D.; Arnarez, C.; Wassenaar, T. A.; Schafer, L. V.; Periole, X.; Tieleman, D. P.; Marrink, S. J. Improved Parameters for the Martini Coarse-Grained Protein Force Field. *Journal of Chemical Theory and Computation* **2013**, *9*, 687–697, PMID: 26589065.

(16) Bereau, T.; Kremer, K. Automated Parametrization of the Coarse-Grained Martini Force Field for Small Organic Molecules. *Journal of Chemical Theory and Computation* **2015**, *11*, 2783–2791, PMID: 26575571.

(17) de Jong, D. H.; Liguori, N.; van den Berg, T.; Arnarez, C.; Periole, X.; Marrink, S. J. Atomistic and Coarse Grain Topologies for the Cofactors Associated with the Photosystem II Core Complex. *The Journal of Physical Chemistry B* **2015**, *119*, 7791–7803, PMID: 26053327.

(18) Periole, X.; Cavalli, M.; Marrink, S.-J.; Ceruso, M. A. Combining an Elastic Network With a Coarse-Grained Molecular Force Field: Structure, Dynamics, and Intermolecular Recognition. *Journal of Chemical Theory and Computation* **2009**, *5*, 2531–2543, PMID: 26616630.

(19) Monticelli, L.; Kandasamy, S. K.; Periole, X.; Larson, R. G.; Tieleman, D. P.; Marrink, S.-J. The MARTINI Coarse-Grained Force Field: Extension to Proteins. *Journal of Chemical Theory and Computation* **2008**, *4*, 819–834, PMID: 26621095.

(20) Marrink, S. J.; de Vries, A. H.; Mark, A. E. Coarse Grained Model for Semiquantitative Lipid Simulations. *The Journal of Physical Chemistry B* **2004**, *108*, 750–760.

(21) Abraham, M. J.; Murtola, T.; Schulz, R.; Pall, S.; Smith, J. C.; Hess, B.; Lindahl, E. GROMACS: High performance molecular simulations through multi-level parallelism from laptops to supercomputers. *SoftwareX* **2015**, *1-2*, 19 – 25.

(22) Kutzner, C.; Pall, S.; Fechner, M.; Esztermann, A.; de Groot, B. L.; Grubmaeller, H. Best bang for your buck: GPU nodes for GROMACS biomolecular simulations. *J.Comput.Chem.* **2015**, *36*, 1990–2008.

(23) Berendsen, H. J. C.; Postma, J. P. M.; van Gunsteren, W. F.; DiNola, A.; Haak, J. R. Molecular dynamics with coupling to an external bath. *The Journal of Chemical Physics* **1984**, *81*, 3684–3690.

(24) Bussi, G.; Donadio, D.; Parrinello, M. Canonical sampling through velocity rescaling. *The Journal of Chemical Physics* **2007**, *126*, 014101.

(25) de Jong, D. H.; Baoukina, S.; Ingolfsson, H. I.; Marrink, S. J. Martini straight: Boosting performance using a shorter cutoff and GPUs. *Computer Physics Communications* **2016**, *199*, 1 – 7.

(26) Bowman, G. R.; Pande, V. S.; Noé, F. An Introduction to Markov State Models and Their Application to Long Timescale Molecular Simulation. Advances in Experimental Medicine and Biology. 2014.

(27) Chodera, J. D.; Noé, F. Markov state models of biomolecular conformational dynamics. *Current Opinion in Structural Biology* **2014**, *25*, 135–144.

(28) Scherer, M. K.; Trendelkamp-Schroer, B.; Paul, F.; Perez-Hernandez, G.; Hoffmann, M.; Plattner, N.; Wehmeyer, C.; Prinz, J.-H.; Noe, F. PyEMMA 2: A Software Package for Estimation, Validation, and Analysis of Markov Models. *Journal of Chemical Theory and Computation* **2015**, *11*, 5525–5542.

(29) Lloyd, S. Least Squares Quantization in PCM. *IEEE Trans. Inf. Theor.* **2006**, *28*, 129–137.

(30) Noé, F.; Wu, H.; Prinz, J. H.; Plattner, N. Projected and hidden Markov models for calculating kinetics and metastable states of complex molecules. *Journal of Chemical Physics* **2013**, *139*.

(31) E., W.; Vanden-Eijnden, E. Towards a Theory of Transition Paths. *Journal of Statistical Physics* **2006**, *123*, 503.

(32) Metzner, P.; Schutte, C.; Vanden-Eijnden, E. Transition Path Theory for Markov Jump Processes. *Multiscale Modeling & Simulation* **2009**, *7*, 1192–1219.

(33) Noé, F.; Schütte, C.; Vanden-Eijnden, E.; Reich, L.; Weikl, T. R. Constructing the equilibrium ensemble of folding pathways from short off-equilibrium simulations. *Proceedings of the National Academy of Sciences* **2009**, *106*, 19011–19016.
